# Consilience in the Peripheral Sensory Adaptation Response

**DOI:** 10.1101/2020.02.17.953448

**Authors:** Willy Wong

**Affiliations:** Department of Electrical and Computer Engineering, and Institute of Biomedical Engineering, University of Toronto, Toronto M5S3G4

## Abstract

Measurements of the peripheral sensory adaptation response were compared to a simple mathematical relationship involving the spontaneous, peak and steady-state activities. This relationship is based on the geometric mean and is found to be obeyed to good approximation in peripheral sensory units showing a sustained response to prolonged stimulation. From an extensive review of past studies, the geometric mean relationship is shown to be independent of modality and is satisfied in a wide range of animal species. The consilience of evidence, from nearly one hundred years of experiments beginning with the work of Edgar Adrian, suggests that this is a fundamental result of neurophysiology.

## 1 Introduction

Consilience is the convergence of evidence from different lines of studies or approaches [54, 55]. The unity of science requires results obtained by one approach to concur with evidence obtained by another approach. This principle has found utility in biology where systematic bodies of evidence can be hard to obtain, and has been invoked most prominently in establishing the modern theory of evolution.

Can consilience find application in sensory physiology? While there is little debate that sensory adaptation appears universally amongst the different senses and organisms, there has been no attempt to carry out a *quantitative comparison* of adaptation responses. Some of the first experiments conducted on sensory nerves were carried out by Nobel Laureate Edgar Adrian. Collaborating with his assistant Yngve Zotterman, Adrian conducted one of his most celebrated experiments: the measurement of rate of impulses from the frog muscle spindle to the stretch of a muscle [1]. What Adrian found was that the neural activity rises immediately upon initiation of stretch and falls monotonically with time. This is now known as sensory adaptation and is observed nearly universally in all of the senses across many different organisms. A schematic representation of their findings can be found in figure 1 which includes spontaneous activity prior to the application of the stimulus (SR), the peak activity that occurs at or soon after the presentation of the stimulus (PR), and the steady-state activity after adaptation has stopped (SS).

**Figure 1:**
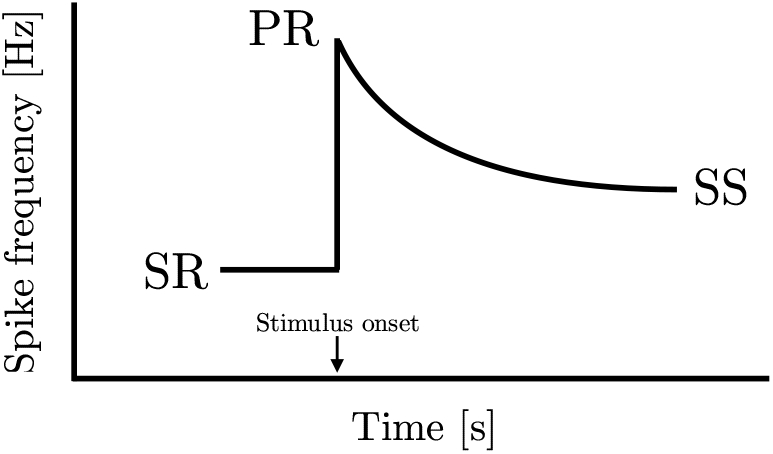
Peripheral sensory adaptation curve. An idealized sensory adaptation response showing steady-state spontaneous rate (SR) prior to introduction of stimulus, the peak response to the stimulus (PR), and the subsequent new steady-state response (SS).

A particular pattern emerges from Adrian’s study when the results are analyzed numerically. By taking the three fixed points in the graph, we observe from his data that the steady-state activity equals the *geometric mean* of the peak activity and spontaneous activity. This finding is not restricted to a single one of his experiments. In the third instalment of the celebrated 1926 papers [2], they measured adaptation in Merkel units in the footpad of a cat to pressure stimuli where a similar result can be found. In equation form, this implies that

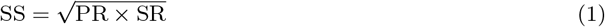

Subsequent to Adrian’s discoveries, many investigators have studied the peripheral response in other modalities and organisms. While the adaptation response tends to follow the same qualitative shape of figure 1, the question that remains unanswered is whether they share *quantitative similarities*. Neurons process and encode many types of information; a diversity of responses can be found at the peripheral level, some of which may even differ from the representation shown in figure 1. Notable exceptions include the rapidly adapting mechanoreceptors in touch as well as certain units involved in temperature coding. This study focuses on responses that show sustained activity with constant input in isolated peripheral sensory units.

An exhaustive search was carried out on past studies of peripheral sensory adaptation. From these studies, results were analyzed and compared to equation (1). Despite vastly different mechanisms and modalities, and in organisms from different phyla in Animalia, equation (1) was found to be obeyed to good approximation. From the perspective of consilience, this demonstrates that equation (1) is widely applicable and may constitute a new law of neurophysiology. The discovery of this equation came from some recent theoretical advances [56].

## 2 Results and Discussion

A comprehensive search was carried out in peripheral sensory adaptation yielding thirty six studies which satisfied the conditions set forth in the search criteria (see Methods). A third of the studies were conducted within the past twenty years. At least fifteen studies included multiple measurements under different conditions. In total, there were 250 adaptation responses analyzed. The dataset spans eight of the most important sensory modalities including proprioception, touch, taste, hearing, vision, smell, electroreception and temperature. A total of thirteen studies were identified to test the same modality/same animal species combination but were conducted in different labs. One study examined units of both high and low spontaneous activity in the same modality. There are other aspects of the data set worth noting and are discussed later.

We begin the analysis with point-wise comparisons to equation (1). Table 1 shows the results of nineteen studies where a single comparison of spontaneous, peak and steady-state activities can be made. Equation (1) appears to hold well across different animal species and modalities, although the limited availability of data in each study makes it difficult to draw robust conclusions. Taken together, however, the convergence of evidence is strong. Error between predicted and measured steady-state response is generally within ten percent or less. There are exceptions. The largest source of discrepancy is found in a study on temperature [39] where in two instances the prediction misses the measured value by a considerable amount. Thermoreception is particularly difficult to reconcile and is discussed in more detail later. A number of results (marked by *) are derived from ‘inverted’ responses and are also discussed later.

**Table 1:** A summary of different peripheral sensory adaptation results highlighting the relationship between spontaneous, peak and steady-state activities for different modalities and organisms. ^*^Indicates data derived from an inverted adaptation response.

| Spontaneous<br>(SR, Hz) | Peak<br>(PR, Hz) | Steady-state<br>(SS, Hz) | Prediction<br>(Hz) | Modality | Organism | Source |
| --- | --- | --- | --- | --- | --- | --- |
| 6.1 | 26.8 | 13.2 | 12.8 | Proprioception | Frog | Figure 10B (exp. 12), [1] |
| 3.7 | 99.7 | 19.3 | 19.2 | Touch | Cat | Figure 8 (exp. 6), [2] |
| 15.4 | 59.9 | 25.9 | 30.4 | Proprioception | Cat | Figure 3 (filled circles, prep. 52), [9] |
| 14.1* | 9.1* | 11.0* | 11.3* | Proprioception | Cat | Figure 3 (open circles, stimulus on, prep. 52), [9] |
| 11.0* | 16.9* | 13.8* | 13.6* | Proprioception | Cat | Figure 3 (open circles, stimulus off, prep. 52), [9] |
| 12.9 | 31.1 | 18.9 | 20.0 | Proprioception | Cat | Figure 4 (initial adaptation, prep. 54), [9] |
| 37.7 | 151 | 81.7 | 75.4 | Hearing | Cat | Figure 6.6 (unit 297-43), [31] |
| 60.1 | 179 | 100 | 104 | Hearing | Cat | Figure 6.7 (unit 299-22), [31] |
| 28.8 | 70.7 | 44.6 | 45.1 | Proprioception | Cat | Figure 7 (first adaptation), [38] |
| 44.6* | 18.1* | 33.4* | 28.4* | Proprioception | Cat | Figure 7 (second adaptation), [38] |
| 33.4 | 65.1 | 43.7 | 46.6 | Proprioception | Cat | Figure 7 (third adaptation), [38] |
| 43.7* | 16.4* | 28.0* | 26.8* | Proprioception | Cat | Figure 7 (fourth adaptation), [38] |
| 28.0 | 57.8 | 42.4 | 40.2 | Proprioception | Cat | Figure 7 (fifth adaptation), [38] |
| 42.4* | 13.4 * | 22.9* | 23.8* | Proprioception | Cat | Figure 7 (sixth adaptation), [38] |
| 23.9 | 43.4 | 32.0 | 32.2 | Proprioception | Cat | Figure 8 (first adaptation), [38] |
| 32.0* | 8.9* | 23.3* | 16.9 * | Proprioception | Cat | Figure 8 (second adaptation), [38] |
| 23.3 | 43.4 | 30.7 | 31.8 | Proprioception | Cat | Figure 8 (third adaptation) [38] |
| 30.7* | 11.2* | 22.9* | 18.5* | Proprioception | Cat | Figure 8 (fourth adaptation), [38] |
| 22.9 | 42.0 | 31.6 | 31.0 | Proprioception | Cat | Figure 8 (fifth adaptation), [38] |
| 31.6* | 7.9* | 23.0* | 15.8* | Proprioception | Cat | Figure 8 (sixth adaptation), [38] |
| 23.0 | 42.0 | 30.9 | 31.1 | Proprioception | Cat | Figure 8 (seventh adaptation), [38] |
| 6.7 | 22.6 | 10.9 | 12.3 | Touch | Cat | Figure 2B (adaptation), [11] |
| 10.9* | 1.9* | 6.7* | 4.6* | Touch | Cat | Figure 2B (recovery), [11] |
| 2.7 | 16.3 | 7.4 | 6.6 | Vision | Squid | Figure 1b, [33] |
| 8.5 | 67.9 | 23.6 | 24.0 | Smell | Mosquito | Figure 2 (top), [12] |
| 11.7 | 69.6 | 30.6 | 28.5 | Smell | Mosquito | Figure 2 (bottom, middle presentation), [12] |
| 112 | 354 | 223 | 199 | Electroreception | Fish | Figure 10 (p-unit, first adaptation), [28] |
| 122 | 359 | 216 | 209 | Electroreception | Fish | Figure 10 (p-unit, second adaptation), [28] |
| 95.6 | 151 | 143 | 120 | Electroreception | Fish | Figure 10 (t-unit), [28] |
| 130 | 639 | 307 | 288 | Electroreception | Fish | Figure 12A, [28] |
| 13.6 | 81.0 | 29.5 | 33.2 | Taste | Rat | Figure 2 (initial adaptation), [49] |
| 4.9 | 27.5 | 11.6 | 12.4 | Proprioception | Crayfish | Figure 1, [3] |
| 12.2 | 96.1 | 28.7 | 34.2 | Hearing (warm) | Gerbil | Figure 1 (upper), [43] |
| 2.6 | 51.1 | 14.5 | 11.5 | Hearing (cool) | Gerbil | Figure 1 (lower), [43] |
| 6.1 | 199 | 30.5 | 34.8 | Smell | Fruit fly | Figure 11A (first presentation), [13] |
| 3.8 | 199 | 28.8 | 27.5 | Smell | Fruit fly | Figure 11C (middle presentation), [13] |
| 1.8 | 9.9 | 4.0 | 4.2 | Proprioception | Cockroach | Figure A i, [44] |
| 3.8 | 199 | 28.8 | 27.5 | Proprioception | Cockroach | Figure A ii, [44] |
| 17.0 | 144 | 33.4 | 49.4 | Temperature | Beetle | Figure 10A, [39] |
| 4.6 | 278 | 21.2 | 35.8 | Temperature | Beetle | Figure 10B, [39] |
| 2.3 | 333 | 26.1 | 27.6 | Temperature | Beetle | Figure 10C, [39] |
| 2.4 | 15.0 | 7.7 | 6.0 | Smell | Fish | Figure 1a (Lys), [20] |
| 4.9 | 70.0 | 22.3 | 18.5 | Smell | Fish | Figure 1c (Trp), [20] |
| 3.1 | 73.2 | 24.2 | 15.1 | Smell | Fish | Figure 1c (Trp, 50 min. later), [20] |
| 7.8 | 61.4 | 19.1 | 21.9 | Temperature | Mosquito | Figure 2A, [23] |
| 3.3* | 15.0* | 7.6* | 7.0* | Temperature | Mosquito | Figure 2B, [23] |
| 0.14* | 2.0* | 0.58* | 0.53* | Vision | Jellyfish | Figure 3D, [21] |
| 1.1 | 2.1 | 1.5 | 1.5 | Vision | Jellyfish | Figure 4D, [21] |
| 1.5 | 40.0 | 7.4 | 7.7 | Water movement | Fish | Figure 2, [41] |
| 5.6 | 104 | 28.9 | 24.1 | Water movement | Fish | Figure 5a, [41] |
| 26.5 | 71.6 | 42.2 | 43.6 | Water movement | Fish | Figure 5b, [41] |

The first two datasets in table 1 are from Adrian’s own pioneering work and are reconstructed using either extrapolated data because the stimulus was not held long-enough for the response to reach steady state [1], or by pooling data from other recordings of the same unit [2]. It would be easy to exclude these studies, but their historical significance cannot be ignored. Of additional interest is a study on hearing which compares the adaptation response of the same unit when cooled or warmed relative to body temperature – see results conducted on gerbil hearing [43]. Changes in local temperature result in spike activities differing by at least a factor of two. One could easily imagine that the three values rise or fall uniformly with a change in temperature. Instead, they move in a direction to preserve the equality of the geometric mean. This has possible implications for the generality of the relationship across different animal species where metabolic activity can differ widely. One study was conducted on the visual sensory structures of the jellyfish (rhopalia) by measuring the output of pacemaker cells [21]. The rhopalia modulates the output of these cells determining the basic swim movement in jellyfish [21, 30].

The unfortunate circumstance is that spontaneous activity is not always reported or shown. However, this can be overcome if the adaptation response is measured to multiple stimulus levels. Since spontaneous activity is the activity in the *absence of stimulation*, its value is independent of intensity. As such, equation (1) predicts a square root relationship between peak and steady-state activity which, on a double-log plot, is a straight line with slope 1/2 and value of intercept dependent on the level of spontaneous activity. See Methods for mathematical details. Figure 2 shows the results from fourteen studies conducted at different levels of intensity. Multiple measurements from the same unit increase the robustness of the findings. In total, this figure comprises over 170 adaptation responses. Panels (a)-(h) show responses from mechanoreception, (i)-(m) chemoreception, (n) thermoreception, and (o) photoreception. Regression analysis is detailed in Methods. From the entirety of the data in figure 2, a single value of slope was found to be 0.662.

**Figure 2:**
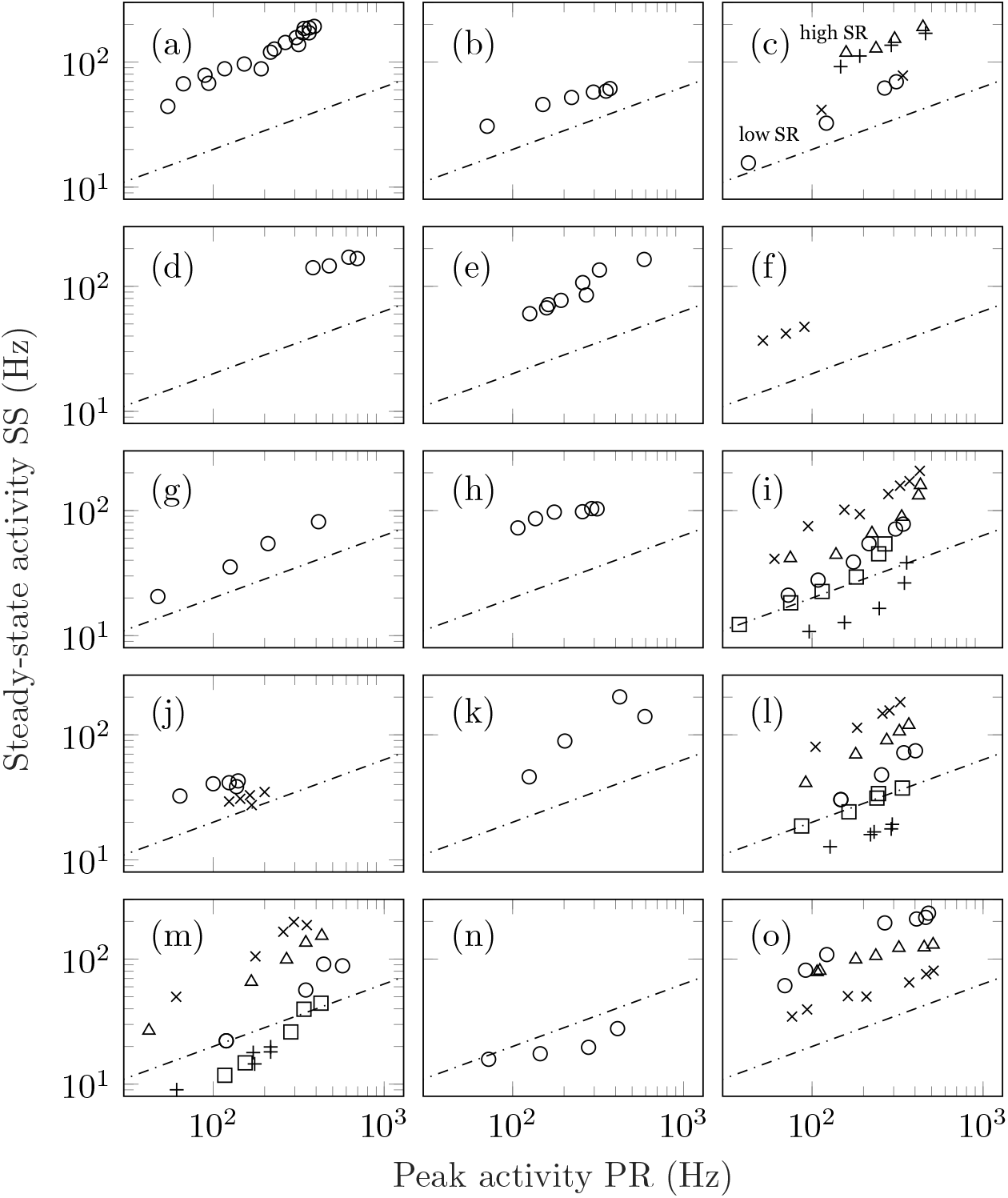
Steady-state activity (SS) plotted as a function of peak activity (PR) for different stimulus intensities. In all panels, the dashed line shows the predictions of equation (1) which is a line with slope one-half on a log-log plot with the value of SR set arbitrarily equal to 4. The actual value of intercept depends on the precise value of the unit’s spontaneous activity. (a) Plot of SS vs PR for auditory data taken from a single guinea pig fibre (figure 1, unit GP-17-4), [50]; (b) the same for auditory data from a single gerbil fibre (figure 1, unit E8F2), [53]; (c) results for four separate guinea pig auditory fibres of both high and low spontaneous activity (triangles, figure 1, unit GP31/08; plusses, figure 2, unit GP27/18; circles, figure 1, GP31/13; crosses, figure 2, GP27/04), [57]; (d) the averaged auditory data from ferrets (figure 6,), [52]; (e) auditory responses obtained from the saccular nerve fibres of a gold fish (figure 3, increment), [18]; (f) responses from lateral line in fish (figure 6), [41] (g) stretch response in crayfish (figures 1 and 2, both PR and SS are shifted upwards by 0.5 log units), [10]; (h) stretch response in frog (figure 3, both PR and SS are shifted upwards by 1 log unit), [34]; (i) response of olfactory receptor neurons in fruit flies (figure 3a: crosses, shifted +0.3 log units; figure 5: triangles, shifted +0.1 log units, methyl butyrate; circles, shifted −0.1 log units, methyl butyrate; squares, shifted −0.3 log units, 1-pentanol; plusses, shifted −0.5 log units, propyl acetate), [37]; (j) taste recordings in fruit fly sensilla (circles, figure 3; crosses, figure 7), [24]; (k) taste response in caterpillar (figure 3), [6]; (l) taste response in blowfly (figure 2a: circles, LiCl, shifted +0.2 log units; triangles, NaCl; crosses, KCl, shifted −0.2 log units; squares, RbCl, shifted −0.4 log units; plusses, CsCl, shifted −0.6 log units), [36]; (m) same as (l) but for figure 2b, [36]; (n) response to cooling in beetles (figure 10), [39]; and (o) vision data from a single ON-centre ganglion cell in the cat. The vision data differ from the other auditory data in that they are derived from pre-adapted luminance values (figure 7: circles, 1 × 10^−5^ cd m^−2^, shifted +0.2 log units; plusses, 1 × 10^−3^ cd m^−2^; crosses: 1 × 10^−1^ cd m^−2^, shifted −0 2 log units), [45].

Of particular interest are panels (a) and (c) which show measurements taken from the same animal/modality (guinea pig hearing) conducted in different labs separated by almost ten years apart. Panel (c) shows the response of four fibres, including two with low spontaneous activity and two with high spontaneous activity. In all cases, the power law relationship is preserved with those units with high spontaneous activity having a higher value of intercept than those with low spontaneous activity. Panels (a)-(d) show mammalian hearing mediated via the cochlea while panel (e) shows hearing in the fish mediated via the otolith organs. In (f), we observe the response of the fish lateral line. Panels (h) and (i) show data from the response of taste receptors in a blowfly to five alkali salts at different intensities. In total, there are forty eight measurements from two units. Although the data are shown with offset, the actual values overlap indicating that they share a similar slope and intercept which is not surprising given the common value of spontaneous activity. The last panel (o) differs from the other studies in that adaptation was conducted *on top* of an existing pedestal. That is, the retinal ganglion cell was adapted to an existing level of luminance before responding to a further increment. Despite the change in test condition, the quantitative aspects of the response remain unchanged.

Adaptation responses measured from ascending and descending staircases allow for further testing of equation (1) to *pre-adapted levels*. A stimulus staircase is a series of ascending and/or descending intensity steps used to probe the response of a unit. Several studies have made use of stimulus staircases including: measurements from warm units in the cat [25] and bat [46] to increasingly warmer temperatures; recordings from M1 cells for non-image-based vision (i.e. intrinsically photosensitive retinal ganglion cells or ipRGC’s) in mice to light of increasing levels [40]; measurements from olfactory sensory neurons of fruit flies to ascending and descending levels of acetone concentrations [32]. The data from four studies together with the predicted values are shown in figure 3. Taken together, this suggests the following generalization of equation (1):

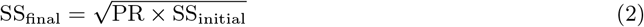

This equation includes (1) as a special case.

**Figure 3:**
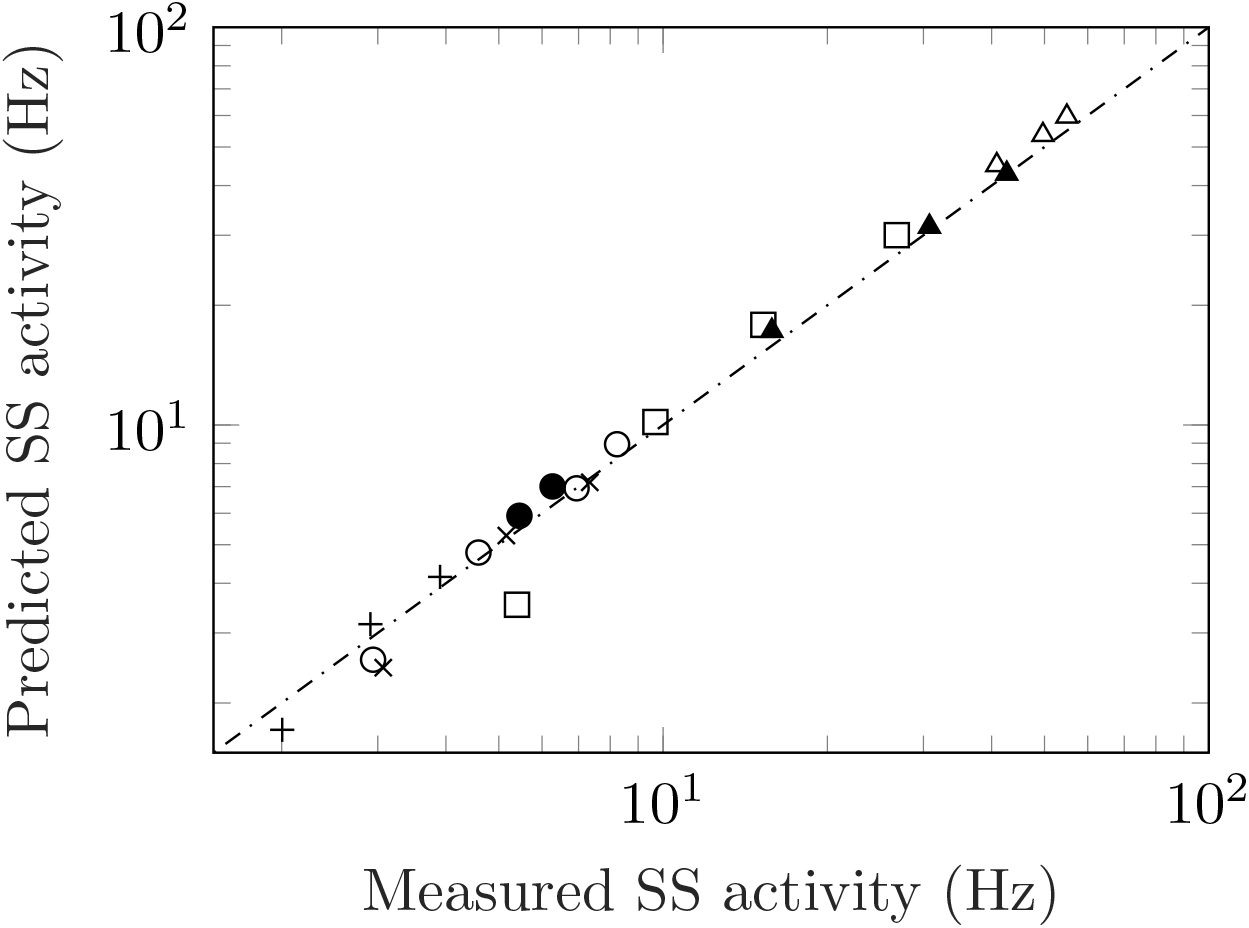
Measured versus predicted steady-state activity for ascending and descending staircases. Data from an ascending staircase of temperatures in warm units of cats (crosses, figure 2a), [25] and bats (squares, figure 2), [46]; response of M1 ipRGC in mice to an ascending/descending luminance staircase (open/filled circles, figures 1b/c), [40]; ascending/descending staircases of acetone concentrations from olfactory sensory neurons in fruit flies which show both regular and inverted responses (open/filled triangles, figure 2c), [32]. Finally, the inverted responses from cold fibres to a descending temperature staircase (plusses, figure 2), [47].

While adaptation responses commonly conform to the schematic representation shown in figure 1, there are also circumstances where the response is inverted from its usual representation. The most common form of an ‘inverted’ response is the recovery after the removal of the stimulus. The spike rate falls before returning to the original value prior to the application of the stimulus. Certain units also exhibit a fully inverted response. These are referred to as *inhibitory responses*, and can be observed in [23] where a warm and cool temperature receptor are presented with the same stimulus: The warm unit follows the typical adaptation response to a rise in temperature while the cool unit falls with the same temperature increase. Inhibitory responses are also commonly found in other modalities. Regardless of its shape, the salient point is that equation (2) is obeyed not just for conventional adaptation but for inverted responses as well. See values marked with * in table 1 and the response to a descending stimulus staircase for odour [32] and temperature [47] in figure 3.

All of this suggests that there is a higher organizational principle underlying peripheral sensory adaptation. What sort of theory or model of transduction would be compatible with equation (2)? There are a number of models, particularly in hearing, that provide good compatibility to experimental data recorded from single unit activity [51, 58]. However, since the geometric mean relationship appears to hold across different modalities, it is not reasonable to expect a modality-specific model to work with other modalities without additional, possibly ad hoc, assumptions. The model of spike frequency adaptation by [5] is likely compatible with the geometric mean relationship provided that a suitable form of the firing rate function is assumed together with an appropriate choice of slope for both the peak and steady-state growth functions. Instead, equation (1) emerges naturally and was first predicted from a theory of sensory processing under development for the past fifty years. The derivation of equation (1) is provided in Appendix 1 and has been detailed fully elsewhere [56].

Few data were found to be in complete violation of equation (1). Those that were are most commonly found in thermoreception. Not only are the responses predicted poorly in table 1 (see results for the beetle), but the slope of the data in panel (n) of figure 2 falls short of the predicted value of 0.5 [39]. In the same study, there are cases where the three fixed points (spontaneous, peak and steady-state) cannot be easily identified from the adaptation response. Other temperature studies show non-monotonic behaviour with changing stimulus levels [46, 47, 26]. While warming responses tend to fit better than cooling, on the balance thermoreception appears to violate equation (1). A small number of results from other modalities were also found to be problematic. One study concerns the taste of blowflies [14]; however, the data in this case was obtained by averaging the response of different sensilla resulting in a non-monotonic adaptation curve. Another study, again concerning taste, shows data for sucrose and salt satisfying equation (1), but not for pheromones [8]. Apart from temperature, however, entire studies found in violation of equation (1) were few in numbers.

Much of this investigation has focused on units that adapt slowly and show a sustained response to continued stimulation. Phasic receptors are rapidly adapting units that do not conform to the representation in figure 1. At steady-state the spike activity is zero. It is believed that phasic units respond to the rate of change of stimulus [7]. A test can be carried out to see if phasic units obey the geometric mean relationship by presenting a time-varying stimulus to induce a sustained response. This was attempted in [19] where the vestibular units of monkeys were subjected to centrifugal forces and an adaptation response was measured to a constant force stimulus. While two of the adaptation responses compare favourably to the equation (unit 213-28: SS_*meas*_ = 80.3, SS_*pred*_ = 86.8; unit 206-18: SS_*meas*_ = 125, SS_*pred*_ = 130), the paper also cites data from units which show far less levels of adaptation (unit 213-28: SS_*meas*_ = 97.0, SS_*pred*_ = 67.6). Moreover, the geometric mean relationship is found only to be satisfied in the activities of *isolated* sensory units in the absence of interaction from other cells in the neural circuitry. Any neurons that are part of the ascending auditory or visual pathway clearly do not follow the geometric mean relationship, e.g. see the auditory interneuron response [27] or the responses of the H1 neuron in the visual cortex [35]. Even within retinal ganglion cells, if the visual signal overlaps both the centre and surrounding regions (thereby recruiting inhibition), this will facilitate responses which deviate from equation (1), e.g. [17].

The main result of this paper is a relationship connecting three fixed points in the adaptation curve. No consideration was made of the time-course of adaptation. And yet, an important point of discussion in the literature is the rate at which adaptation occurs. Adaptation curves are often fitted to a sum of exponentials with different time constants, e.g. [43]. More recently, it has been proposed that neuronal adaptation can be better modelled using a power-law function of time [15]. This raises the question whether equation (1) is compatible with such a formulation. It is important to remember that in both an exponential and a power-law description, a constant offset is required. This offset accounts for the non-zero activity when the stimulus is applied over long periods of time, e.g. see implementation of power law dynamics in [58]. As such, equation (1) is not affected by the time-course of adaptation.

There is considerable debate over the origins and role of spontaneous activity [29]. Spontaneous activity is often thought of as noise within the nervous system when in fact it is clear from equation (1) that it is likely an integral part of normal sensory function. Spontaneous levels can convey information in the same manner that the peak and steady-state levels convey information about the environment [16]. There has been much effort towards investigating the origins of spontaneous activity. Although mechanisms can differ, the geometric mean relationship suggest that the functional role is the same in all modalities. Equation (1) thus appears to imply that all neurons have non-zero spontaneous activity. While this may seem to contradict observation, it is important to remember that there is a difference between low spontaneous rates and zero activity. Assuming a Poisson model of spike generation, the probability that there are no spikes observed in a unit time interval equals exp (−*λ*) where *λ* is the average rate. If *λ* is sufficiently small, spike events will be rare enough to be considered absent even though it is technically non-zero.

While the analysis in this paper has involved the review of a great number of publications, it is virtually impossible to capture all studies that have included measurements of an adaptation response. And yet, the convergence of evidence already is striking. More than 200 measurements taken from different branches of sensory physiology are shown to be compatible with a single equation. These studies span different experimental preparations, different methods of stimulation, and may even require different techniques to measure the response. That the results would conform, even approximately, to the same mathematical relationship is remarkable and illustrates the true nature of consilience. A number of investigators have also engaged in testing the same modality-species combination. Since they were conducted independently, and are shown to obey the same relationship, this speaks to the reliability of the methodologies used. Table 2 shows a summary organized by animal species.

**Table 2:** Summary of studies analyzed organized by animal classification.

| Phylum | Organism | Modality and Source |
| --- | --- | --- |
| Chordata | Bat | Temperature [46] |
|  | Cat | Touch [2, 11],<br>proprioception [9, 38], hearing [31],<br>vision [45], temperature [25] |
|  | Ferret | Hearing [52] |
|  | Fish | Electroreception [28], hearing [18], smell [20],<br>movement [41] |
|  | Frog | Proprioception [1, 34] |
|  | Gerbil | Hearing [53, 43] |
|  | Guinea pig | Hearing [50, 57] |
|  | Mouse | Non-image based vision [40] |
|  | Pigeon | Temperature [47] |
|  | Rat | Taste [49] |
| Arthropoda | Beetle | Temperature [39] |
|  | Blowfly | Taste [36] |
|  | Caterpillar | Taste [6] |
|  | Cockroach | Proprioception [44] |
|  | Crayfish | Proprioception [10, 3] |
|  | Fruit fly | Smell [13, 32, 37],<br>taste [24] |
|  | Mosquito | Smell [12], temperature [23] |
| Mollusca | Squid | Vision [33] |
| Cnidaria | Jellyfish | Vision [21] |

The compilation of studies in this paper also documents the historical development of the sensory sciences which followed the changes in technology unfolding in the twentieth century. For example, the proprioceptive and touch senses were among the first to be investigated as they were the easiest to access and their spike activity slow enough so that reliable spike counts could be achieved even back in Adrian’s time using vacuum tube amplifiers [22]. With the advent of computers and greater access to more invasive regions, the study of hearing and vision with their higher firing rates soon became possible. Finally, taste, olfaction and temperature came relatively later due to the difficulty in controlling and maintaining level of stimulation. This may be one reason why the mechanoreception response shows a higher degree of conformity to equation (1) than chemoreception where the variability can be higher. Nevertheless, it is fascinating to observe that the data recorded in 1926 by Adrian and Zottermann hold almost the same fidelity as modern recordings.

Quoting Paul Willis: “Consilience means to use several different lines of inquiry that converge on the same or similar conclusions. The more independent investigations you have that reach the same result, the more confidence you can have that the conclusion is correct. Moreover, if one independent investigation produces a result that is at odds with the consilience of several other investigations, that is an indication that the error is probably in the methods of the adherent investigation, not in the conclusions of the consilience.”^1^ This work comprises a study of enormous breadth showing, perhaps for the first time, commonalities that exists across almost all sensory modalities and animal species. Evidence includes nearly 100 years of data from eight major sensory modalities, derived from organisms from four major phyla in Animalia (see table 2). Regardless of mechanism or modality, or from which time period the study was conducted, the consilience of evidence lends proof to a new fundamental result of neurophysiology.

## 3 Methods

### 3.1 Data selection and processing

A search of peripheral sensory adaptation studies was conducted in academic databases (Google Scholar, Web of Science and PubMed) casting a wide net using various combinations of keywords including *sense, sensory, adapt, adaptation, fibre, unit, receptor, neuron, afferent, tonic, peripheral, action potential, impulse, spike, inter-spike, interval, ISI, frequency, firing, rate, discharge, activity, PST, PSTH, PETH, peri-event, post-stimulus, histogram, spontaneous, coding, pedestal, staircase, recovery, coding*.

Organism-specific terms like *sensillum* or *sensilla* were also used, as were modality specific terms like *mechanoreception, chemoreception, thermoreception, electroreception and photoreception*. Studies were also found through the tracking of citations. The following inclusion criteria were used:

1. Measurements conducted on *peripheral sensory neurons*
2. Unit stimulated with natural stimuli within its normal sensitivity range
3. Stimulus onset is near instantaneous; stimulus is of sufficient length to achieve a steady-state response
4. Spontaneous, peak and steady-state responses are reported; otherwise, if no spontaneous activity is provided, adaptation response is measured to multiple stimulus levels

In certain cases, restrictions were relaxed to allow a greater number of studies to be included.

Data were digitized and extracted from original publications. The extracted data is available in the Supplementary Data. For certain studies, additional steps were required. For [1], the stimulus was not held long enough to achieve steady-state. Hence, the data from the peak until the removal of stimulus were fitted to an exponential plus offset equation: *c*_1_ exp [−*c*_2_ (*t* − *t*′)] + *c*_3_ where *t*′ is the location of the peak and *c*_1_, *c*_2_ and *c*_3_ are unknown parameters to be determined by a non-linear fitting procedure carried out in MATLAB R2020a (MathWorks) using the function `nlinfit`. From here the value of *c*_3_ was inferred to be the steady-state activity. For [2], the steady-state value was estimated using another experiment conducted on the same unit, but with a slower stimulus ramp (crosses from Fig. 8 of Exp. 6). Other studies have shown that the steady-state activity does not depend on the speed of the ramp, e.g. see [9]. In [34], the 6% stretch was not included as the steady-state value was not provided. For [10], the peak values were obtained from figure 1, but the steady-state values were obtained from figure 2. In [57], only the adaptation values conducted at 5 and 10 dB were extracted for unit GP27/04 as both the 15 and 20 dB experiments show peak responses exceeding the limit of the graph (as noted by the study authors themselves). In [46], the spontaneous activity was missing and only three of four staircase levels were included. For [39], many of the responses do not show clear values for spontaneous, peak and steady-state activities. As such, only the data of figure 10 was considered. For [21], almost all eyes show responses which conform to equation (1). However some responses took a long time to reach steady-state, and thus only the data from the lower lens eye (figure 3) and upper lens eye (figure 4) were included. For [37], several of the responses overlapped making it difficult to track their exact values particularly for the lower intensities. In [40], values were extracted only where there was a clear steady-state level of activity attained, and that the input level lies within the sensitivity range of the unit. This included the third to seventh levels of the luminance staircase in Fig. 1b and the fourth to sixth levels in Fig. 1c.

It is also instructive to examine why certain datasets could not be included in the analysis. One study used large bin widths to calculate firing rates thereby obscuring the fine structure in adaptation [4]. Large bin widths reduce the noisiness of the response at the expense of reducing the value of peak activity. Recent methods have been developed to optimize the choice of bin width for time-varying rate data, e.g. [48]. This method is based on the observation that the spike count per bin accumulated over many trials will converge towards a Poisson distribution. By minimizing the mean integrated square error, an optimal choice of bin width can be found to best estimate the true spike rate. However, use of this method involves taking the entire spike sequence and analyzing it into a number of bins of different sizes, solving for the width that yields minimum estimate error. Many of the adaptation studies cited here were conducted before these methods were available. Fortunately, most studies appear to follow good statistical practice and their results are largely comparable with each other, although the use of adaptive bin sizing would likely improve the estimation of the ‘peak’ in the recovery responses. Some studies have also subtracted away the spontaneous activity from the rate data, e.g. [9], rendering some of the results unusable. Finally, a good number of studies show responses that have not yet reached steady-state before the recording was terminated or the stimulus turned off. This is perhaps the single most significant reason why certain datasets could not be included, e.g. [50, 20]. Or, if they were included e.g. [36, 37], this introduced bias in the analysis. See next discussion.

### 3.2 Regression analysis

An analysis of linear least squares was conducted on the data shown in figure 2. First, a fit was conducted on each dataset separately with the equation

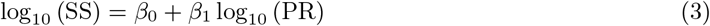

where *β*_1_ is the slope and *β*_0_ the intercept. SS is the steady-state activity and PR the peak activity. If the data conforms to equation (1) then *β*_1_ = 0.5 and *β*_0_ = 0.5 log_10_ (SR), i.e. the base 10 logarithm of spontaneous activity. Since SR is independent of intensity, *β*_0_ is constant. Table 3 shows the results of the analysis carried out in MATLAB R2020a (MathWorks). The analysis shows that much of the variation in the dependent variable is explained by the independent variable (mostly *R*^2^ *>* 0.9). The column labelled ‘Two parameter fit’ in Table 3 shows the values of the fitted parameters together with the root-mean-square error (RMSE). In one dataset, panel c (crosses), there were only two points and thus several of the results are marked as ‘NA’. The confidence interval of the slopes overlapped the predicted value of 0.5 for only 50% of the datasets; there is considerable variation in values. Therefore, it would be difficult to conclude that all datasets share the same slope.

**Table 3:**
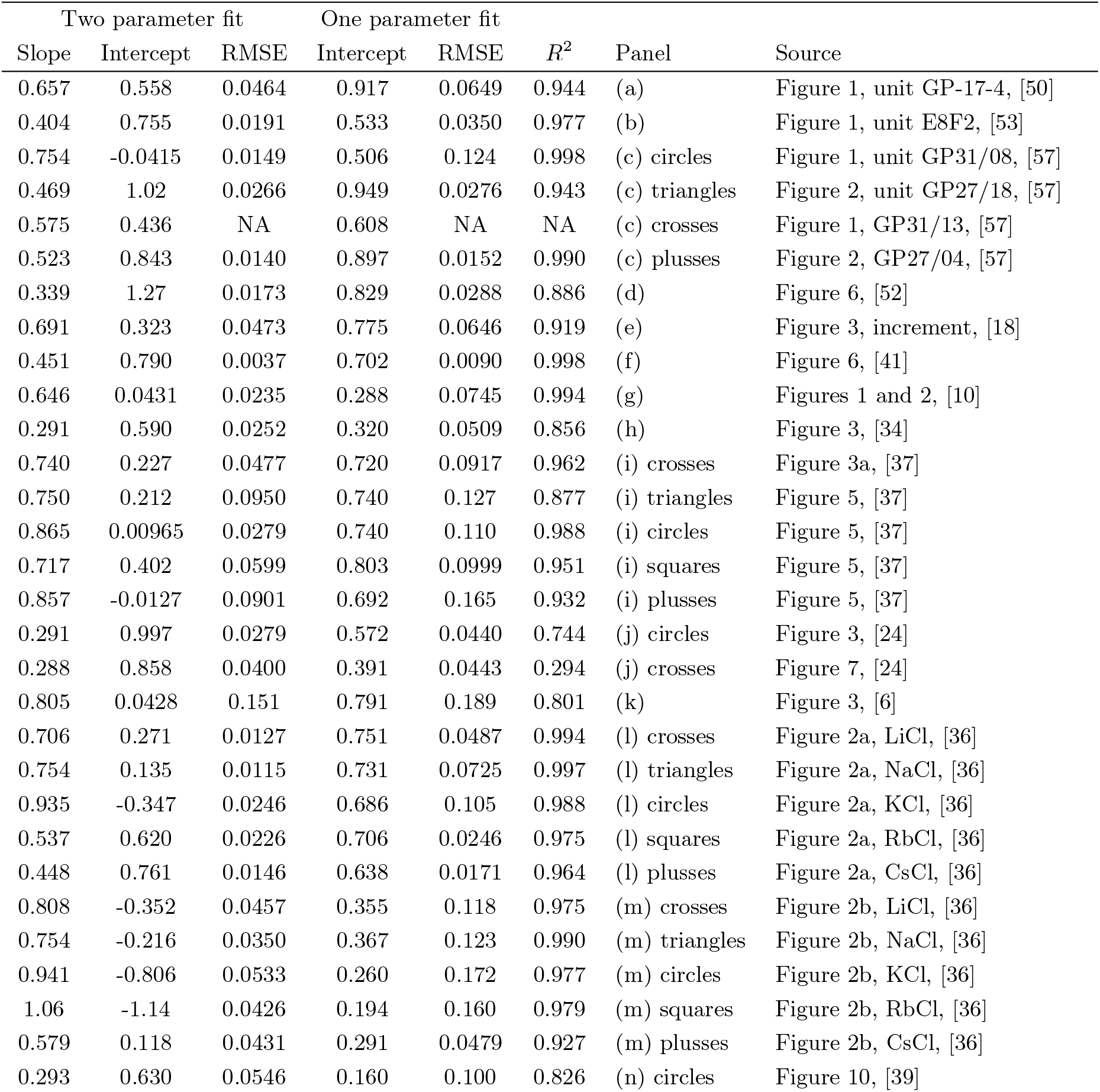
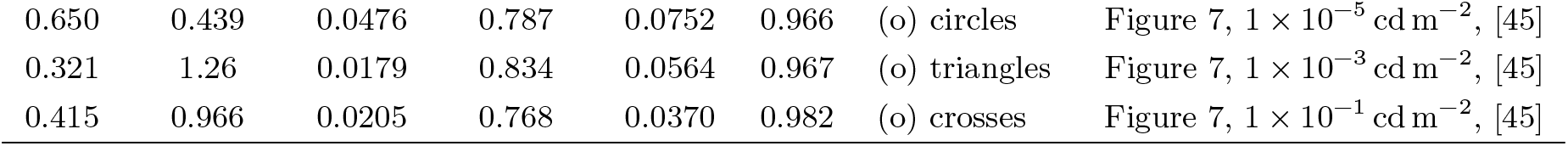
Regression analysis of figure 2.

What are the possible reasons for this discrepancy? Beyond the usual sources of error in measuring spike activity, there is one aspect of the data collection process that has not been addressed. Steady state activity is defined as the activity after which adaptation has stopped. However, as noted in a number of studies, e.g. [57], adaptation curves take longer to reach steady-state when initiated with larger intensities. The consequence of this is significant. Since adaptation is measured from fixed duration presentations, the value of SS obtained from the final portion of the curve may not have reached steady-state, particularly for the higher intensities. This discrepancy will also increase as intensity is increased. From this, we conclude that there can be systematic overestimation of the slope. For example, both [37] and [36] show rate responses which have not yet reached steady-state. This is reflected in the upward rise in points at the higher firing frequencies due to the high steady-state values in panels (i) and (m) in Figure 2.

Nevertheless, if we assume that the slopes for all datasets are in fact identical, we can do one of two things. First is to fit each dataset with a regression line with slope fixed at 0.5. See column labelled ‘One parameter fit’ in Table 3. The second is to fit all of the data to a single value of slope. This was accomplished through a simultaneous curve-fit involving 15 equations with 15 adjustable intercepts and a single value of slope. With all datasets weighted equally, the value of slope = 0.662 was obtained.

## Acknowledgements

The author acknowledges a Discovery Grant (458039) from the Natural Sciences and Engineering Research Council of Canada.

## Appendix 1

### Derivation of Equation (1)

A recent publication detailed the following equations governing the response of peripheral sensory neurons to time-varying stimulation [56]. The theory is based on a mechanism-free approach to sensory information processing:

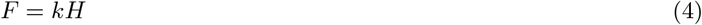

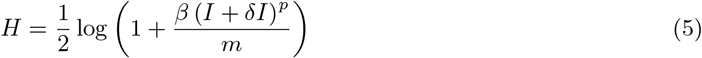

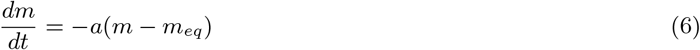

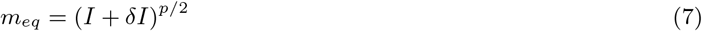

where the firing rate response of the neuron *F* is related to the information or entropy of the stimulus *H* obtained by sampling a signal with intensity *I. m*(*t*) is the sample size and *m*_*eq*_ the optimal value of the sample size. *m*_*eq*_ has dependency on stimulus intensity through equation (7). *k, β, p, a* and *δI* are fixed parameters. The theory has been shown to work well with many time-varying inputs for different sensory modalities and organisms, see also [42]. The equations are not difficult to solve, requiring only a solution to a first-order ordinary differential equation. Moreover, they can be solved even more simply numerically using less than ten lines of computer code.

The following is an abbreviated derivation of equation (1); please see [56] for more details. We begin by solving the response to a step input to obtain the adaptation curve. Given an input that is zero for *t* < 0 and *I* for *t* > 0, equation (6) can be solved to be

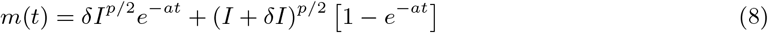

where the continuity of the solution requires the initial condition to be *m*(0) = *δI*^*p/*2^. Substituting *m*(*t*) into *H* and *F* gives the familiar monotonic decay behaviour observed in figure 1.

The simplicity of equation (1) implies that it is likely the result of some approximation. Consider the case where *β* (*I* + *δI*)^*p*^ /*m* ≪ 1 in equation (5). This is satisfied when the parameters are small in value (e.g. *β* ≪ 1) or when the unit is stimulated with lower intensity values, or a combination of both. In this case, we can approximate (5) through a first-order Taylor series expansion. Together with (4), we obtain

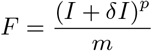

where we have set *kβ* = 2. The values of these constants are not important for this discussion. We are now ready to derive equation (1).

For spontaneous activity, the input intensity is zero and *m* = *δI*^*p/*2^. At stimulus onset, the intensity has value *I* and *m* = *δI*^*p/*2^ through evaluation of equation (8) at *t* = 0. Finally, for steady-state, we evaluate (8) at *t* → ∞ to obtain *m* = (*I* + *δI*)^*p/*2^. This gives

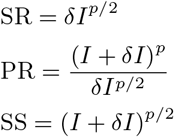

from which we easily obtain equation (1). A similar method can be used to derive equation (2), as well as a corresponding equation for the inverted response.

## Footnotes

1 https://www.abc.net.au/radionational/programs/scienceshow/consilience-powers-the-big-scientific-ideas/5111610

